# Prevalence and genetic characterisation of *Giardia duodenalis* in river water and riverbed sediment using next-generation sequencing

**DOI:** 10.1101/2022.09.01.506148

**Authors:** Muofhe Grace Mphephu, Maggy Ndombo Benteke Momba

## Abstract

*Giardia duodenalis* has been reported in different sources such as water, human stools, animal stools, vegetable farms and markets and soil of public places. However, different assemblages of *G.duodenalis* harboured in riverbed sediments have not yet been investigated. Thus, in this study, we quantified and genetically characterised *G.duodenalis* harboured in the water column and the riverbed sediment of the Apies River to cross this frontier of the unknown in freshwater sediment. Enumeration of *G.duodenalis* cysts was performed by epifluorescence microscopy observation and quantitative polymerase chain reaction (qPCR). Genetic characterisation was achieved by next-generation sequencing (NGS) using the β-giardin gene and bioinformatics analysis of the NGS data. Results obtained through epifluorescence microscopy revealed a prevalence rate of 87% (140/160) of *G.duodenalis* cysts in river water, which was higher than that observed in riverbed sediments (78%, 125/160). However, the qPCR assay showed that gene copies of *G.duodenalis,* which ranged between3.27 log_10_ and 7.26 log_10_ copies/L in re-suspended riverbed sediments, and between 0.49 log_10_ and 3.95 log_10_ copies/L in river water Genetic characterisation revealed six and seven assemblages in river water (A, B, C, D, E and F) and riverbed sediment (A, B, C, D, E, F and G), respectively. Both matrices carried similar sub-assemblages belonging to assemblages A (AI, AII and AIII) and B (BI, BII, BIII, BIV and BV), whereas riverbed sediment carried an additional sub-assemblage BX belonging to the assemblage B. The present genetic characterisation results suggest that Apies River water and its bed sediment harbour considerable quantities of *G.duodenalis* cysts that may cause infections in humans and animals if ingested. Consequently, monitoring of both the water column and respective bed sediments for the presence of *G.duodenalis* is justified to develop strategies for the protection of public health. This study also calls for urgent identification of point sources that are responsible for the contamination of this freshwater source and its sediment.

**Author summary:** 

## Introduction

Water has become increasingly recognised as an effective vector for many outbreaks waterborne diseases in both economically developed and developing countries (1,2,3). Globally, it is estimated that around 2.5 billion people have no access to basic sanitation; in addition, 1.1 billion people worldwide do not have access to safe sources of drinking water, which contributes to the spread of diseases (4,5,6,7). Among causative agents, protozoan organisms have been associated with waterborne diseases and recently appear to have surpassed bacteria, which are considered as the primary concern among waterborne microorganisms (8,9). Protozoan pathogens of animal and human origin have been reported from water sources all over the world (1,2,10,11). Over the past few decades, these pathogens have been increasingly associated with several waterborne disease outbreaks, many of which were associated with public drinking water sources that had met regulatory requirements for turbidity and coliforms (12). In a review of waterborne protozoan disease outbreaks over the period 2017 to 2020 by Ma et al. (2022), 251 waterborne protozoan disease outbreaks were reported during the study period with more such outbreaks reported in developed countries than in developing countries, and the USA and the UK accounting for most of these outbreaks. Europe had the second highest prevalence of waterborne protozoan outbreaks. The United Kingdom, Ireland, Turkey, Poland, Hungary, and Finland have all reported outbreaks of waterborne protozoa. During 2017 to 2020, 49 outbreaks were reported in the UK Asia had the lowest number of outbreaks during this period, but this does not mean that it has done well in case of disease control but indicates the lack of surveillance and the available material for identifying such diseases since most of its countries are still developing stage. These differences in the prevalence of outbreaks between countries can most probably be attributed to the availability of diagnostic capabilities and surveillance programmes to monitor water contamination with pathogenic protozoa (13).Ninety percent of pathogenic protozoan diseases outbreaks reported occur through water, while 10 percent are food related (14).

Among these waterborne protozoa, *Giardia* spp. have been associated with large waterborne outbreaks, thus making *Giardia* the major focus in recent research on waterborne protozoan pathogens (5,7;15). The most notable of *Giardia* spp. is *G.duodenalis* (also known as *G. lamblia* and *G. intestinalis*). *G.duodenalis* is a flagellate protozoan that infects humans and other animals causing a range of symptoms such as diarrhoea, weight loss, abdominal cramps, growth stunting and currently being considered as one of the world’s most common causes of gastroenteritis (16). Giardiasis is considered to be a zoonotic disease because its biological aetiological agent is transmitted by oral-faecal route to humans through animal reservoirs (17). The small dose of infection to produce the disease, together with the small size of the cysts and their environmental resistance allow for *Giardia* dissemination, particularly in vulnerable populations; however, parasitism is present in all countries and at various economic levels (7,18,19). The *G.duodenalis* is transmitted through a thick-walled cyst (8-14 μm). Up to 24 cysts/L have been reported in drinking water (20,21,22), 87 cysts/L in soil (11,23), 0.0087 cysts/L in air (23), and 40 cysts/L in leafy vegetables (Lanata, 2003). *Giardia* cysts have been tracked mainly in water. It is of paramount importance to point out that the prevalence of *Giardia* cysts (cysts/L) in riverbed sediment is yet to be revealed.

Although it is a well-studied organism, known as being responsible for diarrheal diseases in many countries of the world, the presence of *G.duodenalis* in water sources remains unregulated in developing countries However in developed countries, various countries have established guideline values or standards for enteric protozoa in drinking water; the US EPA requires drinking water systems to achieve a 3 log removal or inactivation of *Giardia*; and in Canada the health-based treatment goal for *Giardia* and *Cryptosporidium* is a 3 log (99.9%) removal and/or inactivation of oo(cysts). According to the Surface Water Treatment Rule (SWTR) of the US EPA, public water systems must filter and disinfect surface water and groundwater directly impacted by surface water; 99% of *Giardia* must be removed or killed (https://www.epa.gov/sites/default/files/2015-10/documents/giardia-factsheet.pdf).The prevalence of giardiasis in developed nations is estimated at approximately 2 to 5 percent and in developing countries at 20 to 30 percent (8). Infections due to *Giardia duodenalis* occur worldwide (24,25,26,27),but it is particularly prevalent in areas with poor sanitary conditions and limited water treatment plants. While the Global Enteric Multicenter Study (GEMS) stated that *Giardia* was not significantly and positively associated with moderate to extreme diarrhoea in children under five years of age in sub-Saharan Africa (28,), Feng and Xiao (2011) reported high *Giardia* infection rates in developing countries, including those in Africa. It is estimated that *G.duodenalis* causes 280 million cases of intestinal disease in humans per year, worldwide (10). Research is ongoing due to the continuing public health threat posed by *Giardia* spp.

The development of sophisticated tools and techniques such as next-generation sequencing, whole-genome sequencing and digital PCR has revolutionised the world of microbiology (26,30,31). These new approaches have allowed for a deeper understanding of microbial communities, genetic make-up and enumeration at previously unimaginably minute scales. Although mainly used for bacteria, these techniques have now widely found their way into the identification and characterisation of protozoa such as *Giardia* and *Cryptosporidium* (32,33). Detection of *Giardia* in water is complicated by the fact that the cyst concentrations in water are characteristically low (34). Various types of molecular techniques, such as PCR-based assays, gene cloning and sequencing of different gene types, including housekeeping genes, have proved to be more useful tools due to their high sensitivity and their ability to distinguish between particular assemblages and genotypes, which have led to a greater understanding of molecular epidemiology (1,2,35).

Genes encoding β-giardin (*bg*), triosephosphate isomerase (*tpi*), small subunit ribosomal ribonucleic acid (*SSU rRNA*) and glutamate dehydrogenase (*gdh*) are commonly used for genotyping of *G. duodenalis* isolates. *Giardia duodenalis* organisms are classified into eight genetically distinct genotypes or assemblages (designated as A-H) (36,37,38). Assemblages C to H have been shown to be relatively host-specific, while assemblages A and B are basically unrestricted in terms of the host they can infect (39). Transmission from human to human and animals to human (zoonotic transmission) is orchestrated by assemblages A and B (17). Being host-specific, assemblages C to H cause infections in different types of animals (40). Most studies show that assemblages A and B have been found in clinical samples and are linked to social and economic conditions in their distribution around the world. Multiple infections of both assemblages A and B have been found in different samples, suggesting constant exposure to contaminated sources. Assemblages A and E are commonly found in surface water (41,42,43). In South Africa, *Giardia* cysts have been detected in a variety of waters and treated drinking water and are commonly found in surface waters (44,45). While the importance of the waterborne transmission route of these parasites was recognised 12 years or more ago, waterborne outbreaks continue to occur, and the ingestion of small numbers of *Giardia* cysts is sufficient to trigger an infection. Therefore, in this study, we investigated the prevalence of *G. duodenalis* in Apies River water and its riverbed sediment in order to elucidate different assemblages and sub-assemblages harboured in the above-mentioned matrices.

## RESULTS

### Detection and recovery of *Giardia* using epifluorescence microscopy

Overall, 320 samples collected from the Apies River during the study period consisted of 160 water samples and 160 riverbed sediments. *Giardia* cysts were found in all the examined sampling points of both matrices (16). The number of Giardia cysts recovered in 50 μL of the concentrated original sample ranged between 34 and 118 and between 26 and 60 in riverbed sediment and river water samples, respectively. The observed *Giardia* cysts were captured under the microscope as shown in **Fig S1**. In general, the number of *Giardia* cysts was higher in most riverbed sediment samples than in river water samples, except for sampling sites AR1 and AR12. The observed number of *Giardia* cysts in 50 μL of water was calculated back to 1 L of water (**Table S3**), which was the original volume collected. As illustrated in **Fig 1**, the log10 transformed number of *Giardia* cyst counts in both sediment and river water samples using epifluorescence microscopy ranged from 4.8 log10 to 5.2 log10 cysts per L in re-suspended riverbed sediments, and between3.2 log10 and 5.4 log10 cysts per L in river water. In both matrices, the lowest number of *Giardia* spp. cysts was detected in sampling site AR10, whereas the highest was recorded in AR2. Although the cyst counts were present in high numbers in re-suspended riverbed sediment samples, the calculated prevalence using Equation (1) above revealed a prevalence of 78% (125/160) and 87% (140/160) in sediment and river water samples, respectively **(Table S2).**

**Fig 1:**
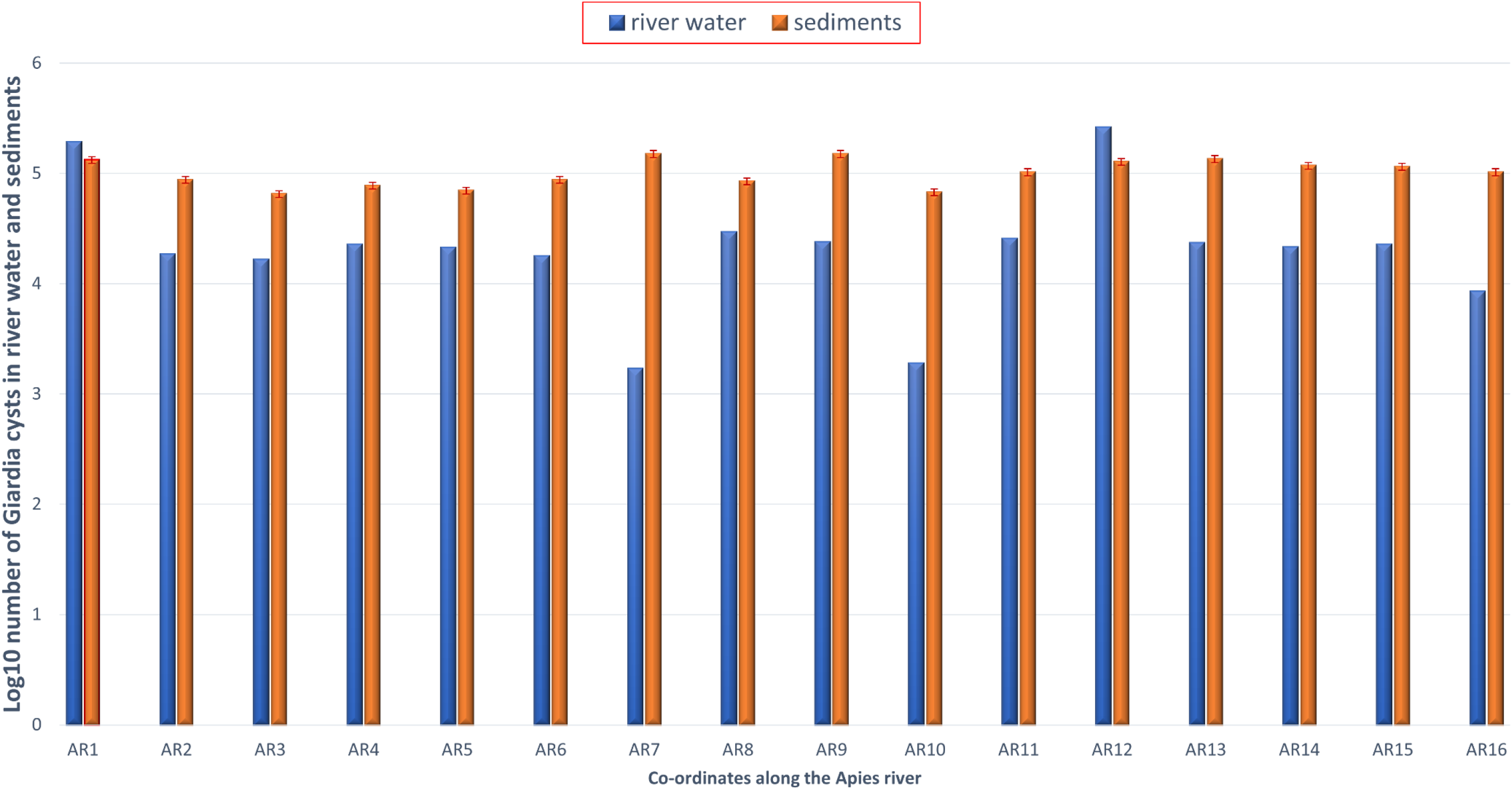
Log_10_ transformed number of *Giardia* cysts in both sediment and river water samples using epifluorescence microsco

### qPCR quantification

Microscopic examination is known to be less sensitive and accurate in terms of the enumeration of microorganisms in environmental samples although it can detect the identity of the target microorganisms. Consequently, quantitative PCR (qPCR), which is one of the PCR-based techniques, was used to complement this method. Nucleic acid-based (NA) detection by PCR is increasingly being used for screening of gastrointestinal parasites, replacing microscopy. The NA-based detection offers greater sensitivity and specificity over microscopic tests and can process more samples in less time. Thus, molecular methods have proven to be useful for the identification and classification of *Giardia* cysts to overcome the limitations of the traditional microscopic procedures. Using the qPCR method, the number of the DNA copies of *Giardia* was determined in both target matrices. The generated standard curve of *Giardia* gBLOCKS targeting the *β-giardin* spanning from 10-1 to 10-8 ng/μL gene copies is shown in Figure S2. This standard curve also showed a strong positive coefficient of determination (R2 = 0.99). The generated Ct values from the PCR machine for both matrices were computed against the standard curve to reveal the number of gene copies of *Giardia* cysts. The amplification curves for both river water and riverbed sediments are shown in **Fig S3-S4**. As shown in **Fig 2**, the number of β-giardin gene copies of *Giardia* cysts obtained using the qPCR assay was higher than the number of *Giardia* cysts observed using microscopic quantification (**Fig. 1**); no amplification was recorded in the river water sample at one sampling point (AR7). The quantification using the qPCR assay is displayed in Figure 2; the log10 transformed number of β-giardin gene copies of *Giardia* cysts ranged between 3.27 log10 and 7.26 log10 copies/L in re-suspended riverbed sediments, and between 0.49 log10 and 3.95 log10 copies/L in river water. The highest number of gene copies per litre was observed in the re-suspended riverbed sediment samples

**Figure 2:**
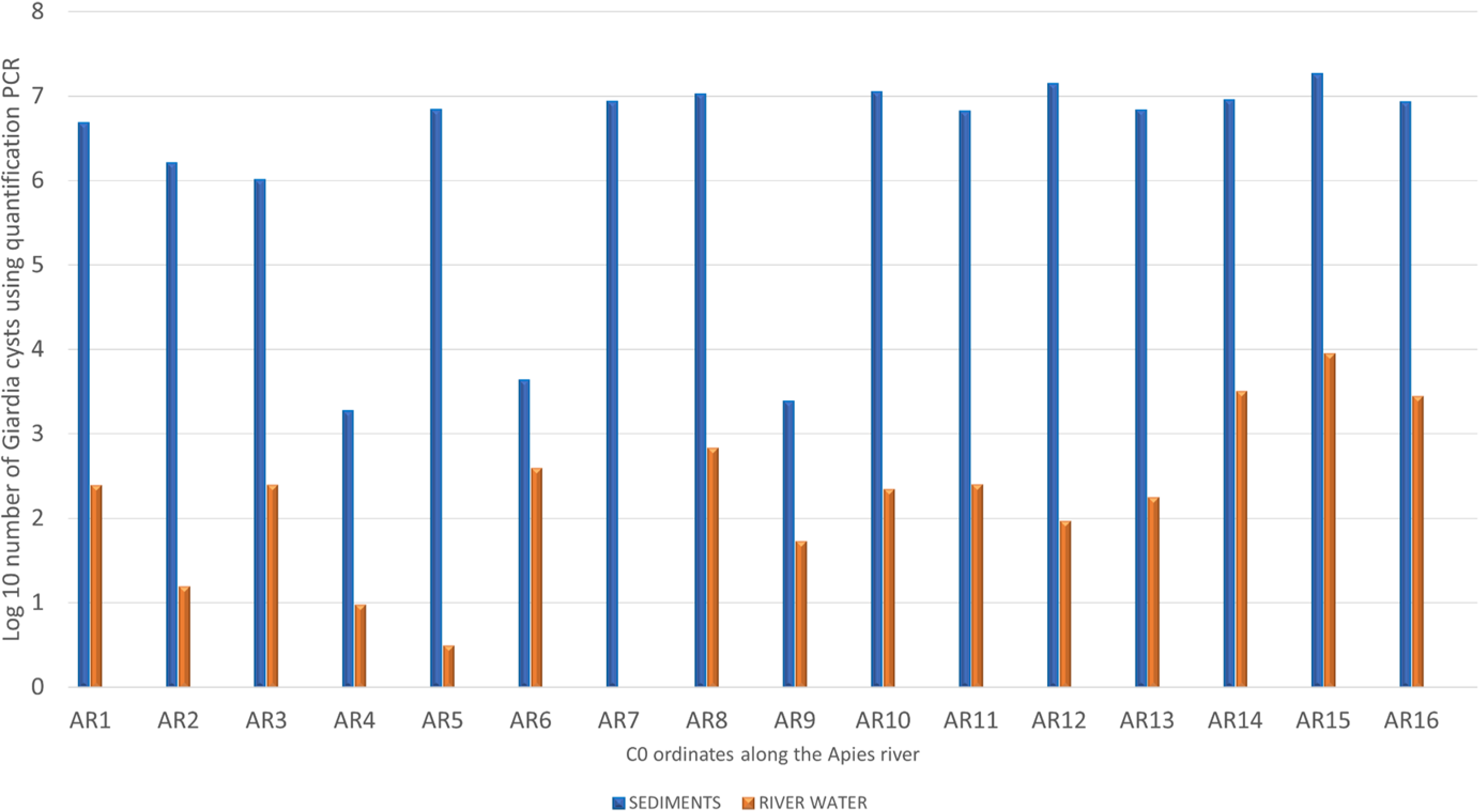
Log_10_ transformed number of β-giardin gene copies of *Giardia* cysts per litre in both sediment and river water samples using qPCR

### Genetic characterisation targeting the β-giardin gene to determine the assemblages and sub-assemblages of *Giardia duodenalis*

Using the nested PCR, the Giardia β-giardin gene was successfully amplified from the genomic DNA extracted from the qPCR positive samples. The PCR products of about 511 bp are shown in **Fig S5**.

Raw reads received from the sequencing centre of the University of Pretoria (South Africa) were separately grouped in one file for all river water samples and another file for all the riverbed sediment samples. These files contained 1 034 780 and 15 384 227 sequences, respectively. After removing the human DNA using DeconSeq V.04.3, the samples were reduced to 960 358 and to 1 372 301 sequences for river water and riverbed sediment, respectively. This process was followed by the removal of chimeric reads and the sequences that did not match the cut-off base pair number. Thereafter, a total of 306 335 high-quality sequences in river water samples and 634 475 sequences in sediment samples were used for the genetic characterisation of *G. duodenalis* as shown in **Table 1**. The assemblages and sub-assemblage sequences from the NCBI (**Table S5)** were used to create a database using the Transeq function in Galaxy software. The generated FASTA files were then used to create a protein database using NCBI BLAST+ makeblastdb (Galaxy tool).

**Table 1:**
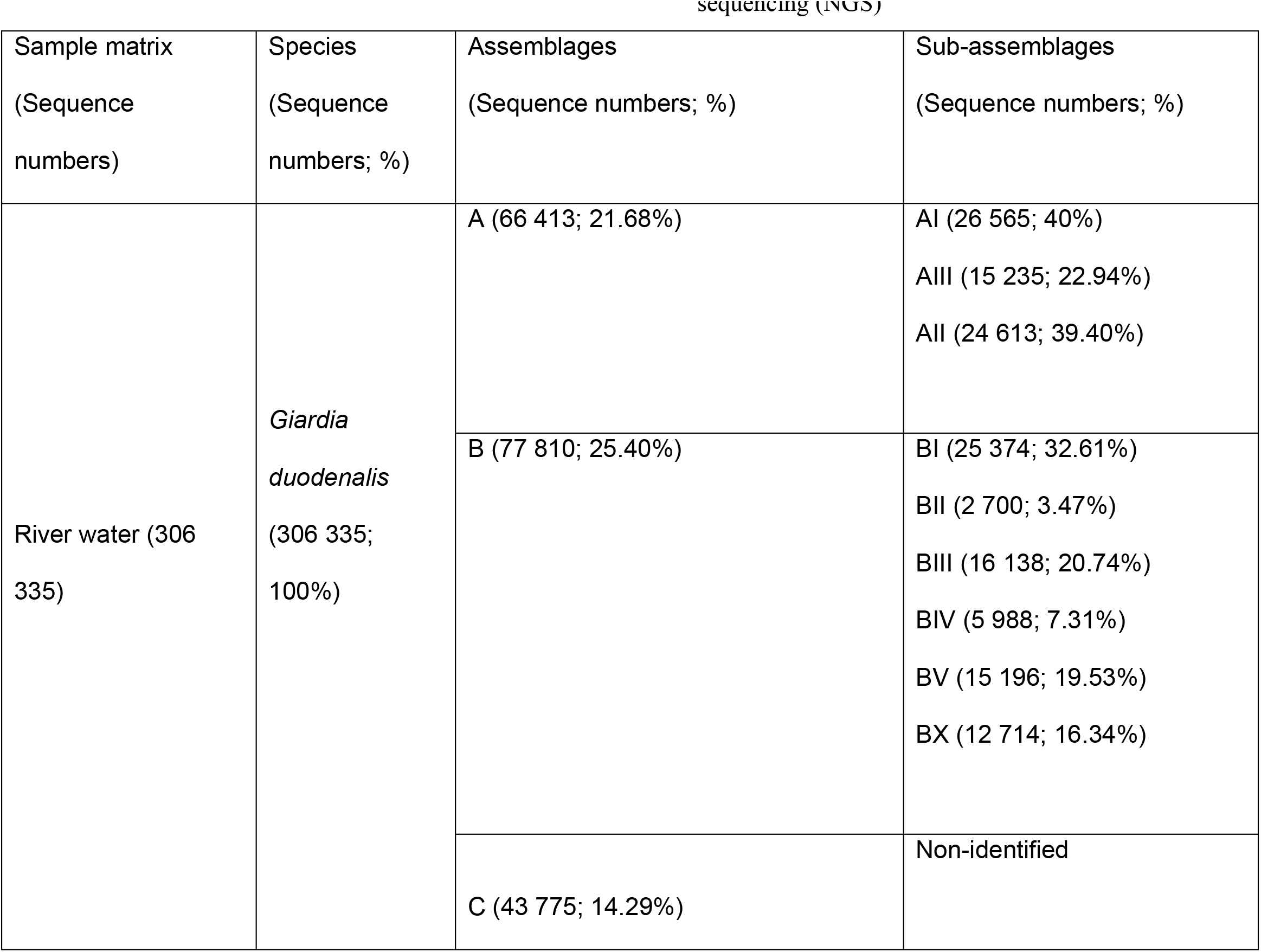
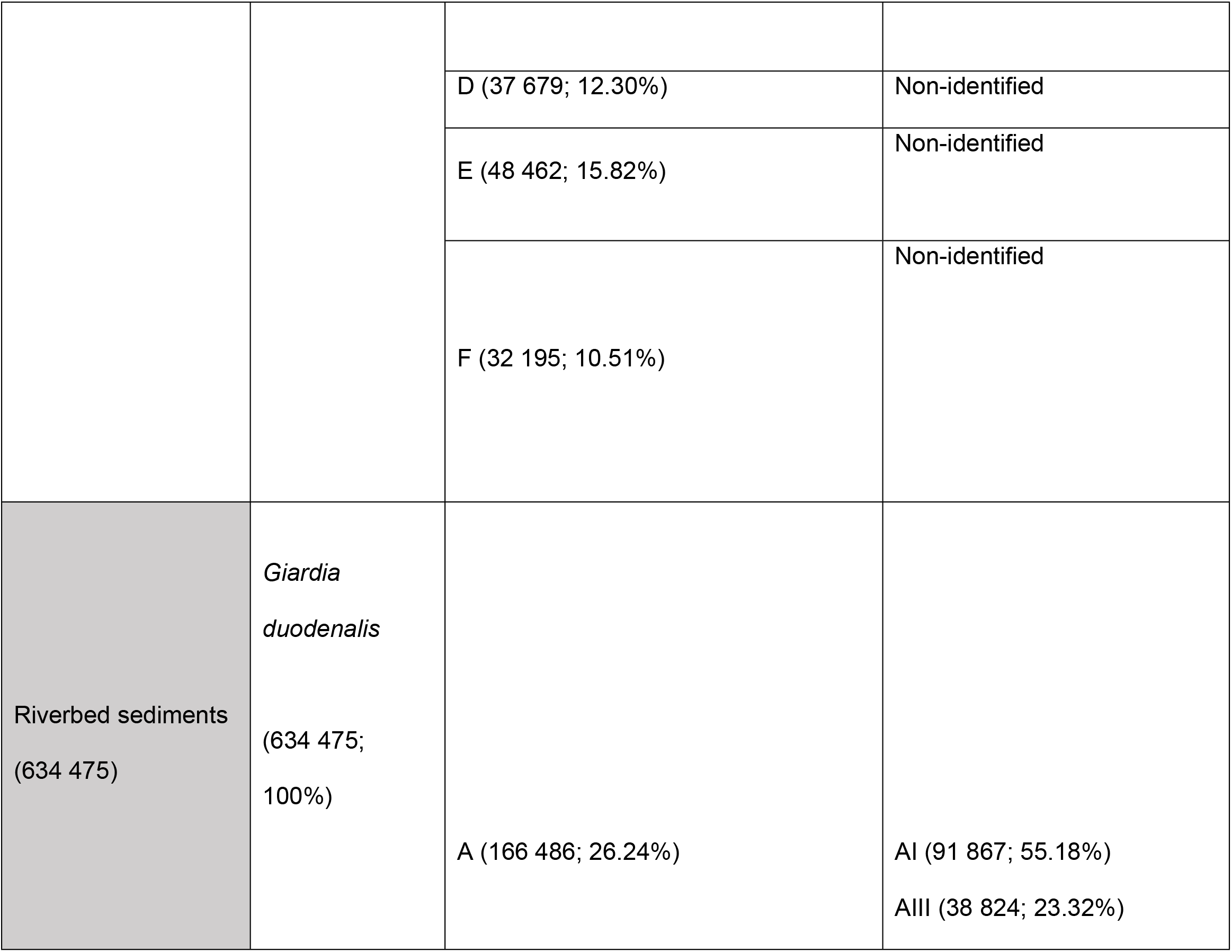
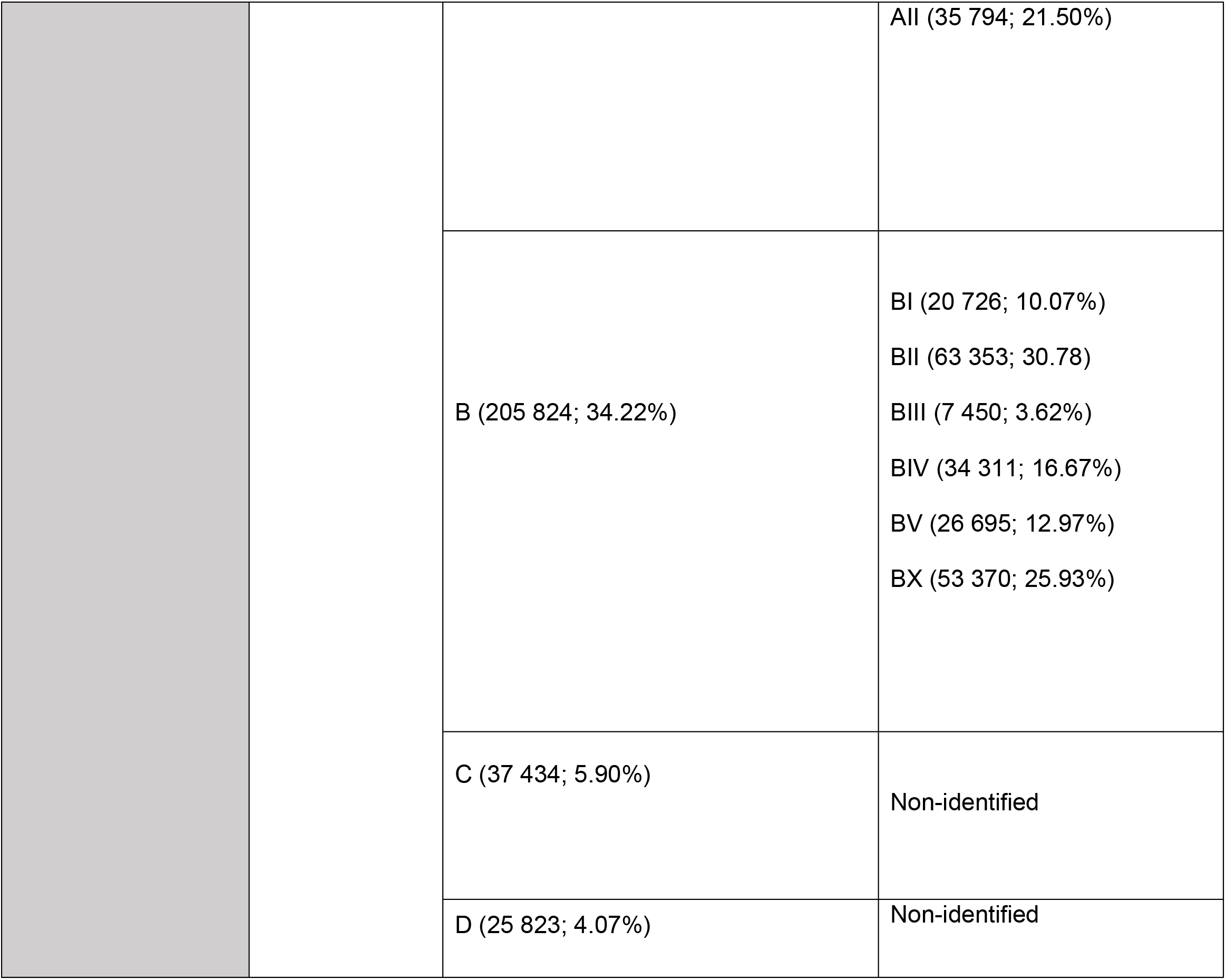
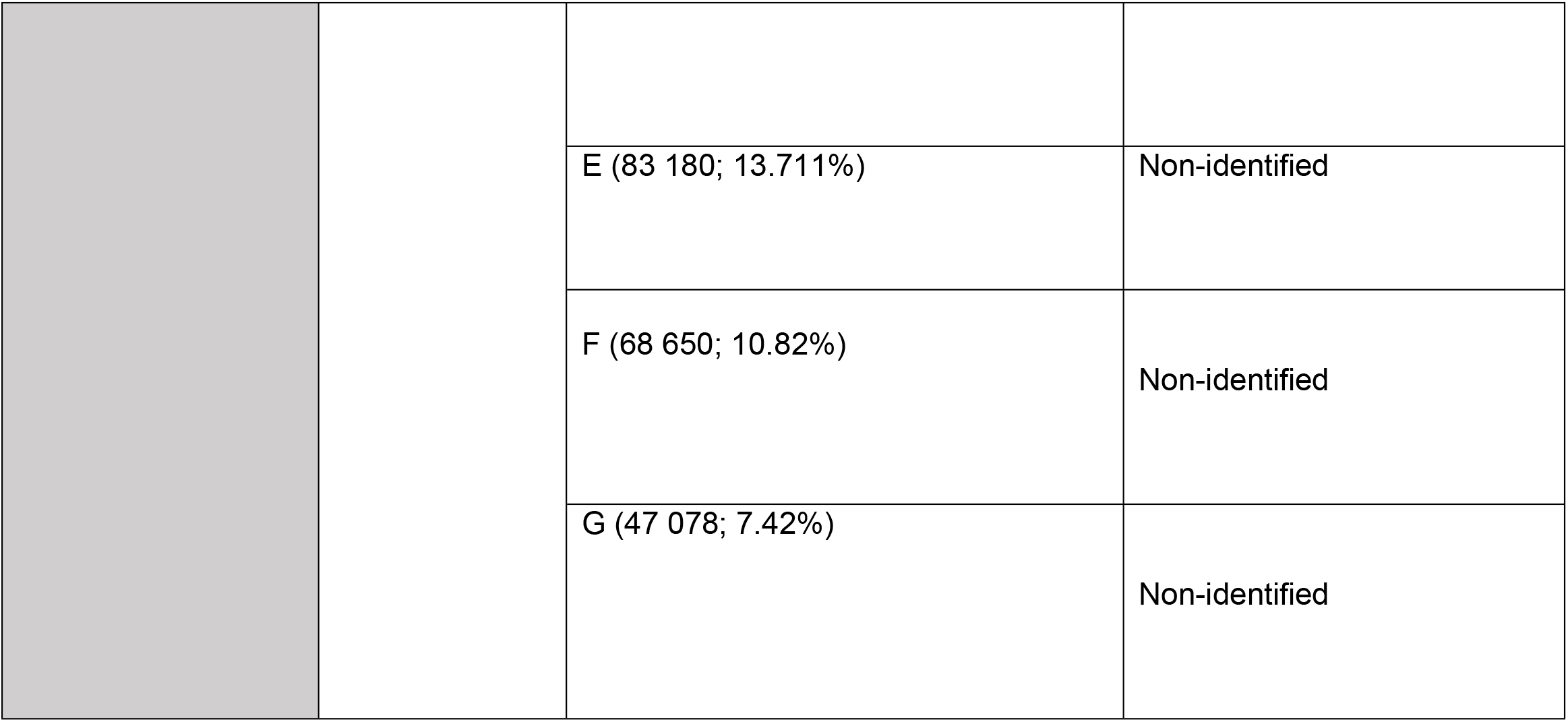
Assemblages and sub-assemblages identified in *G.duodenalis* from two sample types, i.e. riverbed sediments and river water, using next-generation sequencing (NGS)

The assemblages and sub-assemblages of *Giardia* spp. in river water and riverbed samples were identified with next-generation sequencing (NGS) using the *β-giardin* gene. A total of six (6) *Giardia duodenalis* assemblages (A, B, C, D, E, F) were found in river water samples and seven (7) assemblages (A, B, C, D, E, F, G) in riverbed samples. Among the assemblages identified in both river water and sediment samples, assemblage B was found to be the predominant assemblage, followed by assemblage A. The sub-assemblages of only assemblage A (AI, AII, and AIII) and assemblage B (BI, BII, BIII, BIV, BV, and BX) were identified in both river water and sediment samples. The sub-assemblages of the five (5) other *G.duodenalis* assemblages (C, D, E, F, and G) could not be identified. The sequences of the majority of samples were identical to those of five (5) previously reported sub-assemblages, namely B1I and AII in river water samples; the predominant sub-assemblage within assemblage B was BI, followed by BIII, BV, BX, BIV, and BII, while for assemblage A, the predominant sub-assemblage was AI, followed by AII and AIII. In riverbed sediment samples, the predominant sub-assemblage within assemblage A was AI, followed by AIII, and AII; and for assemblage B, the predominant sub-assemblage was BII, followed by BX, BIV, BV, BI, and BIII. The calculated percentage of both the assemblages and sub-assemblages is shown in **Table 4**.

## DISCUSSION

The prevalence of *Giardia* has been well documented in different matrices such as human stools (46,47,48) animal stools (49,50,51), vegetable farms and markets (51,53,), and in soil of public places (52). However, very few studies *Giardia* cysts have been detected in a number of various waters and treated drinking water in South Africa, especially in surface waters. Gerick and co-workers (1995) reported the presence of *Giardia* cysts in raw sewage, treated effluents, surface water and drinking water collected for a period of three (3) years (1992-1994) from former Natal and Transvaal provinces of South Africa. These authors highlighted that all the 10 samples from surface water sites were contaminated with *Giardia* cysts. Mackintosh et al., (2000) also pointed out the presence of *Giardia* cysts for the first time in March 1998, while the Stellenbosch Municipality had been testing quarterly for this protozoan parasite at the three (3) untreated raw water sources. In 2010, Dungeni and Momba reported the presence of *Giardia* spp. in wastewater and treated effluent samples collected between January and April 2008 from 4 wastewater treatments (Refilwe, Baviaanspoort, Zeekoegat and Rayton) of the Gauteng Province of South Africa. As can be seen these few studies have been conducted in South African surface waters, and little has been known especially for the sediments. Furthermore, no data on waterborne giardiasis outbreaks are available among the significant number of waterborne outbreaks reported (56,58,59).

The results of the present study indicate that *Giardia* is highly prevalent in both riverbed sediment and river water of the Apies River. Both matrices were positive for *Giardia* cysts, indicating a relationship between river water and its riverbed sediment. The presence of *Giardia* in Apies River water and sediment is a matter of great concern, given the multiple uses of this water source by the surrounding communities. Moreover, there is also an indication of the possibility of faecal excretion of cysts either by infected animals or humans. This contamination might originate from domestic sewage and drainage of contaminated water from either agricultural practice such as livestock farming or runoff from vegetable fields fertilised with manure contaminated with cysts. Contamination of this water resource, which is used by the population residing along the Apies River and the neighbouring communities, alerts us to the fact that there is a risk of infections. *Giardia* cysts are highly infective, environmentally robust pathogens, ubiquitous in domestic and wild animals, and capable of zoonotic transmission. These factors might have also played a role in its prevalence in the Apies River since adequate sanitation is still lacking in the area and there is a high possibility of wastewater spillage (60). *Giardia* cysts are very resilient and may survive in water for months, increasing the risk of the population to be infected by *Giardia*. Also, they are well known to be resistant to chemical disinfection (61,62), such as chlorination, which has always posed challenges for water treatment authorities, wastewater treatment plant operators and catchment authorities. Although chlorination has been widely used as a disinfectant in water and wastewater treatment processes against many pathogens, it appears not to be very effective in eliminating protozoan pathogens (44).

The Apies River runs along the areas where both rural and urban activities overlap, thus contributing to many sources of contamination. This, in turn, increases the infection rate and populations are at risk since street vendors use the Apies River water to wash fruit and vegetables for sale. Water from the Apies River used for crop irrigation represents a considerable impact on the Tshwane Municipality of South Africa as it has many small-scale agricultural farms. Furthermore, communities around this water source live in close proximity to their companion animals and pets, which are also reservoirs for Giardia cysts. The informal dwellers allow their pets to roam freely, consume garbage food and defecate indiscriminately. It is known that animals infected with *Giardia* such as dogs, cats, livestock, rodents and wild animals may play an important role in *Giardia* transmission, as they can be reservoirs of zoonotic infections (63). Sanitation in developing countries, especially in sub-Saharan Africa, is still a challenging factor for most populations, and therefore microbial load remains a major concern. It is noteworthy that river contamination in some of these areas could be due to possible defecation in the river, or cysts from contaminated land areas being washed away into the river by runoff during heavy rainfall events, resulting in heavy environmental contamination. Human or animal wastes have been implicated as sources of environmental contamination in reported waterborne protozoan disease outbreaks over the years (31,64,65,66). Therefore, it is crucial to monitor the quality of water sources in terms of protozoan parasites.

Microscopic examination can detect the identity but not the potential infectivity of the *Giardia* cysts (Caccio et al., 2003). Thus, molecular methods have proven to be useful for the identification and classification of *Giardia* cysts to overcome the limitations of these traditional procedures. Quantitative PCR has revealed a high quantity of *Giardia* cysts and was able to detect the highest and the lowest DNA concentration. The qPCR assay reduces the time taken and the risk of contamination and offers increased sensitivity (1,2,31,68,69). Molecular characterisation using the β-giardin gene as a target is considered to be unique to protozoan parasites, in that it allows the rapid identification of *Giardia duodenalis* and its assemblages in environmental samples. In the present study, seven (7) of the *Giardia duodenalis* assemblages were identified in sediment samples (A, B, C, D, E, F, G), and six assemblages in river water samples (A, B, C, D, E, and F). However, the sub-assemblages were only identified within assemblage A and B. Among these, three (3) sub-assemblages within assemblages A (AI, AII, and AIII) and six (6) sub-assemblages (BI, BII, BIII, BIV, BV, and BX) within assemblage B were identified in river water and sediment samples (Table 4). Little is known about the prevalence and genetic diversity of *Giardia* assemblages D, F and H, except that they are largely host-specific (30,38).

Results of the present study revealed assemblage B as the main assemblage within the majority of the water samples. This is consistent with other studies that reported assemblage B as the predominant assemblage in water (31,70). Assemblages A and B are associated with human infection; therefore, they pose a serious health risk. As mentioned above, the river water under this investigation is generally used for agricultural irrigation purposes, and therefore it poses a significant risk of infection to humans, through the consumption of raw produce. Assemblage B has been found in vegetables and fruit (71) and some of these vegetables can be eaten raw. This puts farmers and customers at risk because of continuous exposure to the contaminated water through fruit and vegetable processing as the cysts can survive disinfectants and other treatment processes. Furthermore, *Giardia duodenalis* assemblage B has been shown to cause infection with a longer duration of illness compared to cases infected with assemblage A. Nevertheless, it is important to note that both assemblages A and B are associated with zoonotic infection and most attention has been given to assemblage B. Among the six (6) sub-assemblages (BI, BII, BIII, BIV, BV, and BX) identified within assemblage B, it has been reported that sub-assemblage BIII infects both humans and animals, while sub-assemblage BIV appears to be human-specific (72,73). Of the three sub-assemblages of assemblage A, two (AI and AII) are mainly found in human infections and AII is occasionally found in livestock. Sub-assemblage AI is far more prevalent in animals and wildlife; this is considered as a major sub-assemblage of zoonotic concern. Our study identified six sub-assemblages of assemblage B, with BIII and BIV being commonly isolated from African populations, and this reflects the complex circulation of the parasite in the environment.

The sub-assemblage E was found to be at 6.42% and 10.47% in our study and is mostly associated with livestock such as sheep and dairy cattle worldwide (74,75,76,77). Livestock using surface water as a drinking water source may contract *Giardia* from contaminated water. Adult sheep produce approximately 1-3 kg of faeces per day, leading to environmental and possibly surface water contamination (78). Once an infected host defecates, cysts in the faeces can be transported to surface water by runoff, typically from rainfall events (79,80). *Giardia* can cause significant economic losses for sheep farmers. This parasite has been associated with scouring (diarrhoea) and reduced carcass weight in lambs. Scouring is a significant problem as it predisposes sheep to cutaneous blowfly myiasis, and this is associated with increased costs for producers and health risks for sheep. Scouring also results in increased microbial contamination of carcasses and reduces the efficiency of meat processing. Sheep have been found to act as a source of zoonotic infection and recently sub-assemblage E has been linked to human infection as an emerging human pathogen. This, therefore, increases the risk of exposure of humans to infections. Farmers in neighbouring communities along the Apies River have been known to farm sheep and a lot of work is required for the production of wool and meat, and thus the workers are constantly in contact with the animals putting them at risk of infection. Animal studies have found that concentrations of *Giardia* cysts in sheep may reach > 504 cysts/g (81). With this high cyst concentration excreted it is inevitable that at some point farmers and their workers will become infected. Dairy cattle are more at risk and it is known that dairy cattle generate a good income for farmers due to the high risk of infection of calves which act as the primary source of infection. The high prevalence of *Giardia* in both Apies River water and its sediments has many implications such as serious health risks for consumers of this water, the loss of income for farmers, and the potential of the river to act as a transmission route for zoonotic assemblages of *Giardia.*

Based on the findings of this study, it is concluded that the Apies River water and its sediments are contaminated with *Giardia duodenalis* and therefore this source poses serious health risks to consumers of the water. The significant proportion of the isolates examined in this research indicates that the river water and sediment are heavily contaminated. The possible *Giardia* transmission route presented by this source of water necessitates immediate action to reduce the potential health impact of waterborne diseases in the community. An adequate sanitary survey of this water source is strongly advised, as well as the implementation of water and sanitation projects in the community. Most of the assemblages and sub-assemblages identified in the present study are frequently responsible for human infection, which therefore supports the need for better sanitation and hygiene practices. The molecular characterisation using the β-giardin gene has revealed the distribution of assemblages of *Giardia* in aquatic environmental settings such as river water and its bed sediments. Consequently, monitoring of both the river water and its respective bed sediments for the presence of *G.duodenalis* is justified to develop strategies for the protection of public health. This study also calls for urgent identification of point sources that are responsible for the contamination of this water source and its sediment.

## Materials and methods

### Site description, sample collection and preparation

The full description of the study sites, sample collection and preparation has been presented in detail by Mphephu et al. (2021). Briefly, river water and riverbed sediment samples were collected from July to December 2018 along the Apies River in Gauteng Province, South Africa at 16 different coordinates (AR1-AR16). Sediment samples were processed as described by Abia and co-workers (2015). Sediment supernatant and water samples were concentrated by filtration through the Pall Envirochek™ 1 μm HV capsule filter (Pall South Africa (Pty) Ltd., Midrand, South Africa) using a vacuum pump at a pressure of 200 kPa. Concentrates in the filter were recovered in 125 mL of elution buffer containing Laureth-12 (PPG Industries, Gurnee, IL, USA), 1 M Tris buffer pH 7.4, EDTA and silicone antifoam agent (Thermo Fisher Scientific, USA), and visualised as described below.

### Microscopic enumeration of *Giardia* cysts

A direct smear was prepared. A drop of about 50 μL of the water sample was placed on the slide and allowed to cover the surface. The wet mount was examined firstly for mobile parasites at 100X magnification. After this examination, a drop (approximately 10 μL) of Lugol’s iodine (diluted in a 1:5 ratio with sterile deionised water) was added to the edge of the slide, and a cover slip was placed over the slide. The slide was examined for *Giardia* cyst structures at 40X magnification. The prevalence rate of *Giardia* cysts (counts) was calculated using Equation (1) as follows:

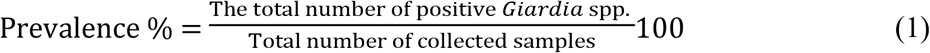

### DNA extraction

For river water samples, DNA extraction was carried out using ZymoBIOMICS™ DNA Microprep Kit (Catalog No. D4301 & D4305) following the manufacturer’s instructions with slight modifications, as described by Mphephu et al. (2021). For sediment samples, the DNA was extracted using freeze and thaw method as described by Nichols and co-workers (2006) also with a few modifications as described by Mphephu et al. (2021). After DNA extraction, the removed DNA for both water column and sediment samples were quantified using the NanoDrop 2000 Spectrophotometer (Thermo Fisher Scientific) and stored at −20 °C until further examination.

### Quantitative PCR

To quantify absolute gene copies through the qPCR assay, a standard curve converting the cycle threshold (*C*_t_) values to *Giardia* spp. gene copy numbers was constructed by using the serially diluted gBLOCKS® Gene Fragment covering the *β-giardin* gene (IDT, WhiteSci, South Africa) as a template in an optimised qPCR assay. The full sequence of gBLOCKS Gene Fragment covering the *β-giardin* gene is depicted in **Table 2**. A graph of *C*_t_ values versus log_10_ (gene copy number) afforded linear calibration curves with typical R^2^ values of 0.991, efficiency of 94.7% and slope of −3.472

**Table 2:**
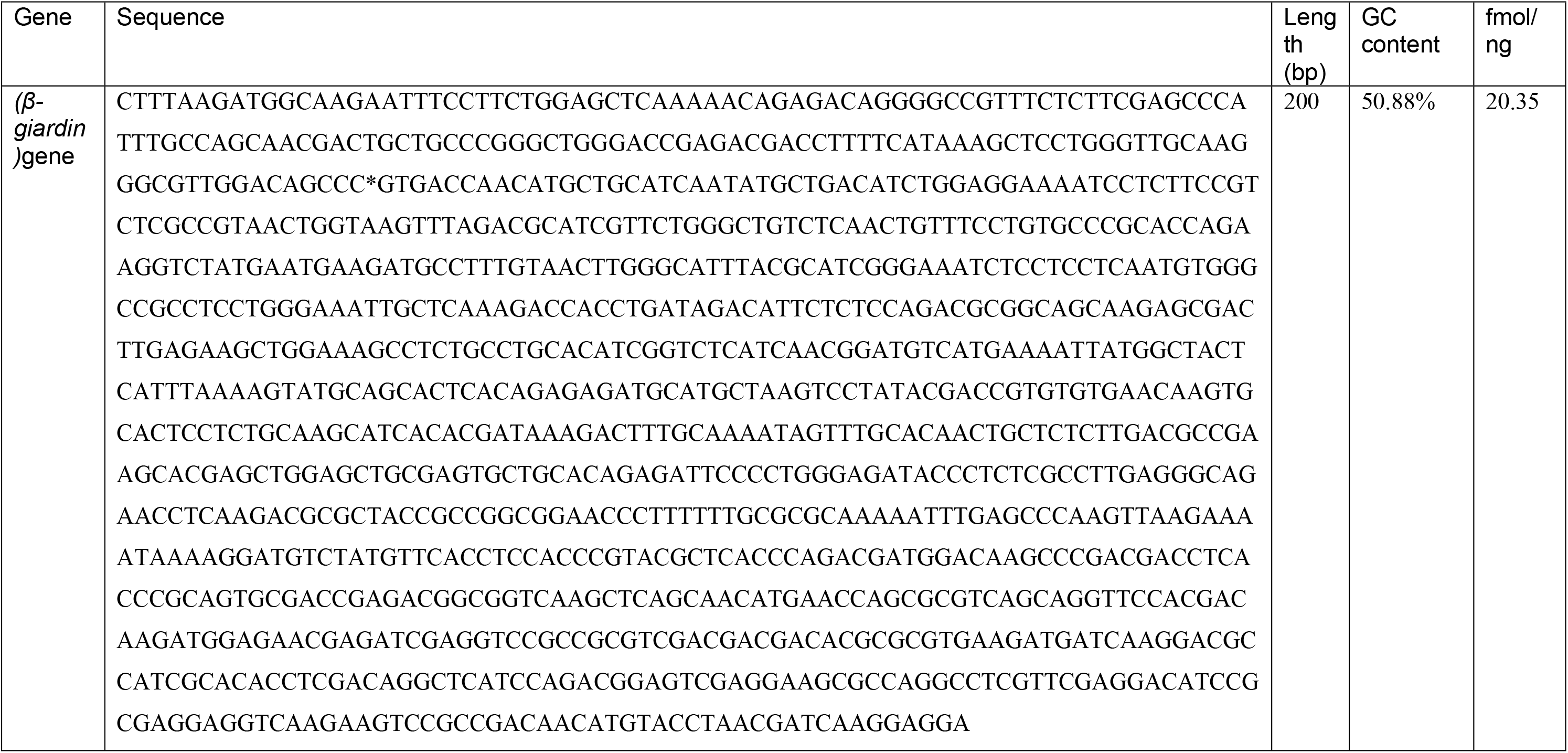
gBLOCKS oligomer for internal control and standard curve generation

The qPCR assays were used to determine the gene copy numbers of *Giardia* spp. in both river water and riverbed sediment samples. Amplification reactions were conducted in a total volume of 20 μL containing 5 μL of template DNA, 10 μL of the PowerUp™ SYBR™ Green Master Mix (Thermo Fisher Scientific, South Africa), 0.25 μL of each forward and reverse primers (Tongesayi and Tongesayi, 2017) (synthesised by Inqaba Biotechnical Industries (Pty) Ltd., South Africa), listed in **Table 3,** and 4.5 μL of nuclease-free water (Thermo Fisher Scientific, South Africa). The purified DNA of *Cryptosporidium* spp. (negative) and of *Giardia* spp. positive control both from Microbiologics, Thermofisher, South Africa was included in each PCR run as controls. Cycle threshold (*C*_t_) values were calculated by the CFX96 Touch™ Real-Time (Bio-Rad, South Africa). The qPCR reaction was run in five steps as follows: enzyme activation at 50 °C for 2 min (step 1); hold Dual-Lock™ DNA polymerase at 95 °C for 2 min (step 2); denaturation at 95 °C for 15 s (step 3); annealing at 95 °C for 15 s (step 4); and extension at 55 °C for 1 min (step 5). The last three steps were repeated 40 times. The amplification curves were observed and captured on the CFX96 Touch™ Real-Time PCR Detection System (Bio-Rad, South Africa).

**Table 3:**
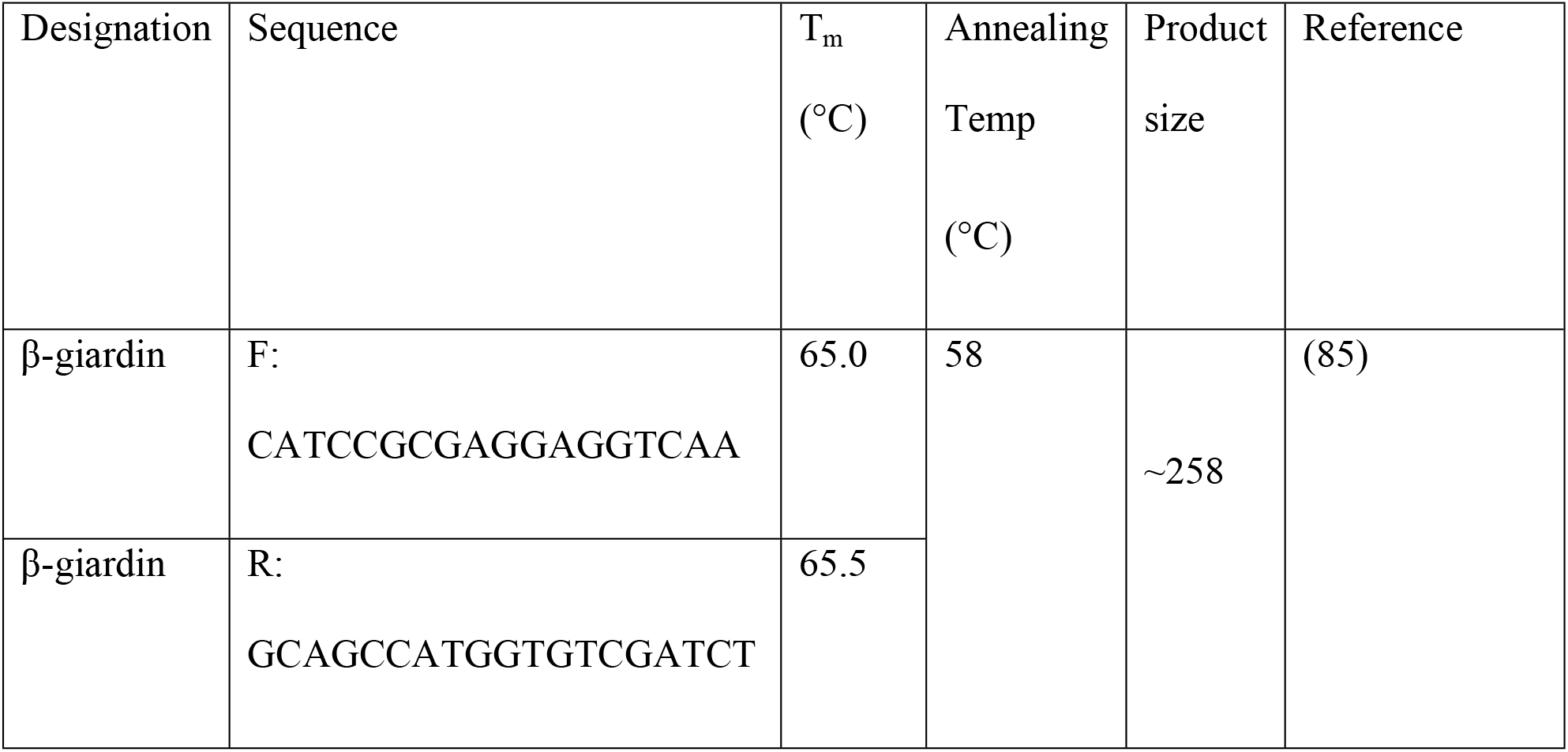
Oligonucleotide primers used in qPCR assay.

**Table 4:**
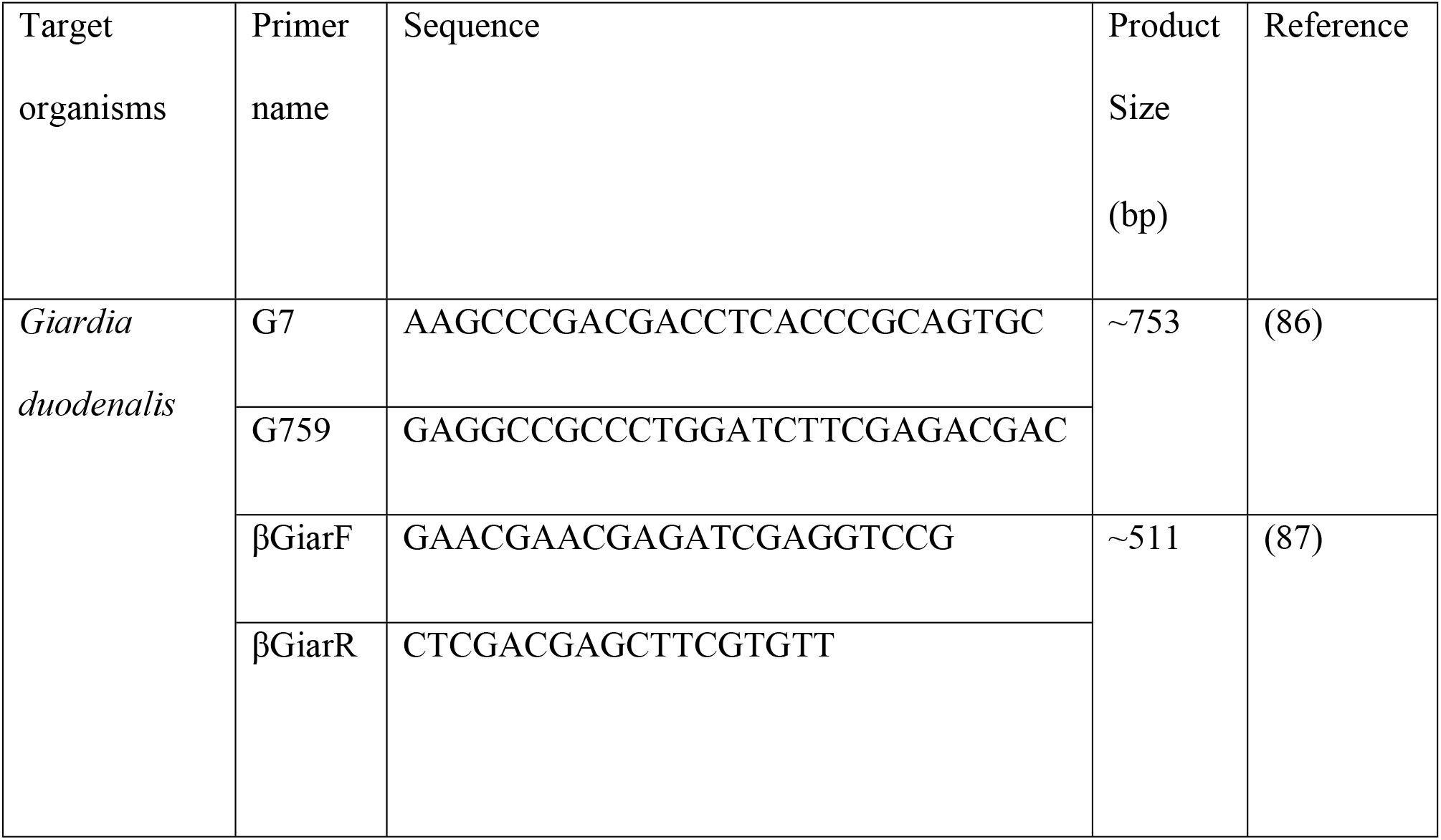
*Giardia* spp. primers used for nested PCR.

### Nested PCR, DNA sequencing and bioinformatics analysis

For the genetic characterisation of *Giardia* spp., nested PCR amplification of the *β-giardin* gene was conducted on a SimpliAmp™ Thermal Cycler (Thermo Fisher Scientific, South Africa), using the primers depicted in **Table 4**. A 605 bp fragment was targeted for the first round and a 532 bp fragment was targeted for the second round of the nested PCR assay. The PCR reaction was run in a total volume of 25 μL consisting of 2.5 μL of 10×buffer containing 1.5 mM MgCl_2_, 0.5 μL of 200 mM of each dNTP, 0.5 μL of 10 pmol of each primer, 0.125 μL of ExTaq DNA polymerase and 5 μL of purified DNA and nuclease free water to 25 μL. All the reagents were purchased from Promega (South Africa). The nested PCR assay was performed under the following cycling conditions: after an initial denaturation step of 5 min at 94 °C, a set of 35 cycles was run, each consisting of 30 s at 90 °C, 30 s at 65 °C and 60 s at 72 °C, followed by a final extension of 7 min at 72 °C. A positive (*Giardia* spp. DNA), negative (*Cryptosporidium* spp. DNA) template and non-template controls were included in each PCR run. After amplification, the DNA fragments from the secondary PCR reaction for *Giardia* spp. were separated according to their size by gel electrophoresis, which consisted of 1% (w/v) agarose gel (Inqaba Biotechnical Industries (Pty) Ltd., South Africa) in Tris acetate buffer (40 mM Tris-HCl, 20 mM EDTA at pH 7). A middle range DNA ladder (Thermo Fisher Scientific, South Africa) was used in the first lane of wells as a reference marker. The gels were run at 300V/1000mA/150W in a gel tank (Bio-Rad, USA). Gels were observed on the GelDoc Imaging System with the Syngene Gel Documentation system (Vacutec (Pty) Ltd., South Africa). The resulting amplicons of the desired base pair (bp) range were purified using the PureLink™ PCR Purification Kit (Thermo Fisher Scientific) following the manufacturer’s instructions. Amplicon sequencing was performed as described by (82). Briefly, amplicons were shipped to Stellenbosch University where they were sequenced using the MiSeq platform (Illumina, USA) following manufacturer’s instructions. Generated raw reads were subjected to bioinformatics analysis as described below.

The primers listed in **Table 4** were used to process the generated reads to keep only reads that corresponded perfectly with the primers (no mismatch permitted) using FastQC v.0.72+galaxy1 software (88). DeconSeq v.0.4.3 was used to eliminate human DNA contamination (89). Good quality sequences were further developed in the *de novo* mode for the removal of chimeric reads with UCHIME (90). The high-quality sequence readings were BLASTed against an in-house database of *β-giardin* gene for species and assemblages in the NCBI **(Table S1** given in the Supplementary section).

## Acknowledgements

This research article is a part of a master’s dissertation and received funding from the Department of Science and Technology (DST) and National Research Foundation (NRF) through the South African Research Chairs Initiative (SARChI) Chair for Water Quality and Wastewater Management (grant number: UID87310) and the Tshwane University of Technology. MGM obtained a scholarship from the NRF Freestanding, Innovation, and Scarce Skills Development Fund (grant number: UID17553). Opinions expressed and conclusions drawn in this study are those of the authors and are not necessarily attributed to the funders.

